# Plasma Induced Afterglow Luminescence

**DOI:** 10.64898/2025.12.21.695776

**Authors:** Jinxin Zhang, Xiao Chen, Zhitong Chen, Pengfei Zhang

## Abstract

Afterglow imaging, is an emerging optical modality using agents that emit persistent luminescence after excitation ceases to eliminate tissue auto fluorescence and improve signal-to-background ratios. However, there was limited number of organic matters could produce persistent luminescence when using light, ultrasound or ionizing radiation as the excitation source. Herein, we explored cold atmospheric plasma (CAP) as a universal and exclusive stimulus for activating intense and long-lasting afterglow from widely-exist organic matters, which could not easily excited by light, ultrasound or X-ray radiation. Furthermore, we demonstrated the biomedical applications of the CAP-induced long-lasting afterglow for photodynamic residual removal after surgical resection of tumor. It should be point out that using CAP as the excitation source might opened new avenue for afterglow imaging clinical translation based on some FDA-approval organic matters learning from the drug repurposing strategy.

## Introduction

Luminescent phenomena are a consequence of the energy absorption, at moderate temperature, that occurs in a material when exposed to any exciting source and have motivated ongoing explorations of their better performance in many advanced applications, such as information storage, light-emitting diodes (LEDs), bioimaging and sensing, etc [1-2]. Different types of luminescence can be distinguished according to the nature of the exciting source (photoluminescence, electroluminescence, radioluminescence, mechanoluminescence, sonoluminescence, chemiluminescence, bioluminescence, etc) [3-4]. However, most of them depend on real-time excitation, which could induce autofluorescence from the tissue during biological imaging and greatly reduce the sensitivity and specificity of imaging within living organisms [5].

Persistent luminescence, also known as afterglow, is a phenomenon in which the material shows long-lasting luminescence after the cessation of the excitation source and has motivated ongoing explorations as the candidate of luminescence with real-time excitation manner. Through the continuous effort of researchers, a variety of persistent luminescence materials with appealing properties have been successfully created in both inorganic and organic systems to store photon energy in trap states and delaying light emission for several hours to days after excitation with light, ultrasound, X-rays, etc [6]. Despite advances in technology and enhanced knowledge of luminescence, translation of these materials into clinical applications has been far slower than expected because of the time-consuming and labor-intensive evaluation on their long-term biosafety and metabolic clearance [7]. Learning from the drug repurposing strategy for identifying new uses for approved or investigational drugs that are outside the scope of the original medical indication might be an effective alternative approach to traditional luminescence materials for clinical translations depending on new energy source for persistent luminescence excitation.

Plasma is a state of matter along with solids, liquids and gases. Plasma can be divided into high-temperature, thermal and non-thermal groups [8-9]. As one group of the non-thermal plasma, cold atmospheric plasma (CAP) is a partially ionized gas generated from a portable device at room temperature and atmospheric pressure, comprising a unique cocktail of energetic species (including electrons, ions, reactive oxygen and nitrogen species, and mild UV photons) [10-11]. Cold atmospheric plasma has been emerging and highly promising technology with broad applications in biomedical area including cancer therapy, chronic disease management, antimicrobial applications[12-14]. However, there was little report on using cold atmospheric plasma as energy source for persistent luminescence excitation.

In this study, we report an innovative finding that atmospheric cold plasma has a universal and unique stimulating effect on sustained luminescence. We demonstrate that brief exposure to CAP jets can instantly ignite strong and long-lasting afterglows of various materials, which remain completely dark under prolonged illumination or ultrasonic excitation. This study systematically investigated the exclusivity and universality of this effect across various material categories. We further optimized the afterglow performance by adjusting plasma parameters and conducted preliminary research on its potential mechanism, which seems to be related to the ability of plasma to produce multiple active substances. Our work not only introduces a new platform for activating and studying persistent luminescence, but also greatly expands the range of materials that can exhibit this characteristic, opening up highly potential opportunities for anti-counterfeiting, biological imaging, and intelligent sensing.

## Materials and Methods

### Material Preparation

Materials were purchased from commercial suppliers and used without further purification. The material is dissolved in DMSO to increase their solubility, and then diluted the samples with ultrapure water for subsequent experiments at a concentration of 2.5%.

### Set up apparatus for CAP-induced luminescence imaging

CAP induced luminescence imaging system consists of four parts: (1) CAP generation system includes a power supply, a CAP transducer, a gas source, and a CAP excitation device. The CAP jet device used in this study consists of a metal electrode rod (diameter 2 millimeters) and a grounding electrode (annular copper foil) (Figure S1). (2) Luminous collection system: A cooled charge coupled device (CCD) from IVIS Lumina XR imaging system (Caliper, USA). (3) The imaging darkbox comes from the IVIS Lumina XR imaging system. (4) For the “CAP induced luminescence imaging” mode, place the CAP device in an external imaging dark box. These material solutions are stored in glass bottles or EP tubes, and CAP devices are placed above the solution samples to excite the materials. After CAP excitation, transfer these solution samples to a dark box and collect luminescent images by cooling a CCD camera. The imaging parameters are as follows: bioluminescence mode; Open the filter; Domain View: C.

### For measurement of CAP-induced afterglow emission band

Used for measuring CAP induced luminescence emission bands. The material concentration is pre irradiated with plasma at 50 μg /ml or 2000 μg /ml for 3 minutes. Signals collected through four different channels after excitation: GFP (510-570nm), DsRed (570-650nm), DsiRed (570-65nm), and DsaRed (570-600nm), Cy5.5 (690-770 nm), and ICG (820-880 nm).

### Cell culture and animal model establishment

The mouse cancer cell line 4T1 (ATCC) was cultured in DMEM medium supplemented with 10% fetal bovine serum at 37 °C and 5% CO_2_ atmosphere. To induce tumors, 1×10^^6^ 4T1 cells suspended in 100 μL PBS were subcutaneously injected into 6-8 week old female BALB/c mice. Monitor tumor growth through caliper measurement and make the tumor reach approximately 100 mm^3^ before surgical resection to establish a postoperative recurrence model.

### Surgical procedures and recurrence monitoring

Before tumor resection, 5-ALA drug was orally administered or injected in situ into the tumor, and mice were anesthetized with isoflurane. The primary tumor was removed under sterile conditions, and the remaining tumor tissue was less than 1mm. The wound was sutured with surgical sutures. CAP treated photosensitizer was injected in situ into the tumor site, and mice were kept in the dark. Tumor recurrence and weight measurement were monitored every day.

### Photosensitizer surgical navigation and CAP-induced luminescence imaging in vivo

Through preliminary material screening, we have selected 5-ALA used clinically as photosensitizers. For surgical navigation applications, CAP treated drug induced afterglow luminescence has been developed, displaying extended tumor retention time (>2 hours) to achieve precise afterglow imaging guided surgery. For in vivo imaging, the use of CAP treated drugs through in situ injection enhances tumor localization and tumor edge localization, achieving precise phototherapy with simultaneous resection and treatment. All surgeries were performed under the guidance of fluorescence navigation.

### Statistical Analysis

All data were presented as the mean ± standard deviation (SD) from three independent experiments. Statistical analyses were performed using SPSS 21.0 software (SPSS, IBM, IL, USA). Comparative analyses were performed using a two-tailed Student’s t-test. Statistical significance was set at p < 0.05.

## Results

### Universal and Non-Invasive Activation of Persistent Luminescence by Cold Atmospheric Plasma

To investigate whether cold atmospheric plasma (CAP) could serve as a general stimulus for persistent luminescence. we evaluated 174 compounds. Quantitative bioluminescence imaging revealed that >32% of compounds exhibited detectable afterglow following CAP treatment. We further screened 20 representative compounds, including clinical drugs, fluorescent dyes, and optoelectronic materials, and systematically compared the effects of light, ultrasound, and CAP in the material solution state (Figure 1A-B). White light illumination and ultrasound stimulation failed to elicit appreciable afterglow in the majority of tested compounds under matched excitation doses, consistent with their dependence on specific photophysical or phononic properties. In contrast, brief CAP exposure (≤ 3 min) produced bright, sustained afterglow signals in nearly all tested materials. CAP-induced luminescence intensities were significantly higher than those induced by conventional stimuli (p<0.001, one-way ANOVA with Tukey’s correction), demonstrating that CAP functions as a universal and non-invasive “charging source” capable of overcoming the limitations of conventional excitation modalities (Figure 1C). Notably, we selected 5 compounds that exhibited strong CAP-induced afterglow in solution for subsequent solid-state characterization. The CAP-induced afterglow intensity of these compounds in the solid state was 3–5 times higher than that in the solution state (Figure 1D-E). This observation suggests that the dense and ordered molecular/particle packing in the solid phase minimizes the scattering and loss of CAP-excited energy, thereby facilitating the efficient conversion of excitation energy into afterglow emission.

**Figure 1.**
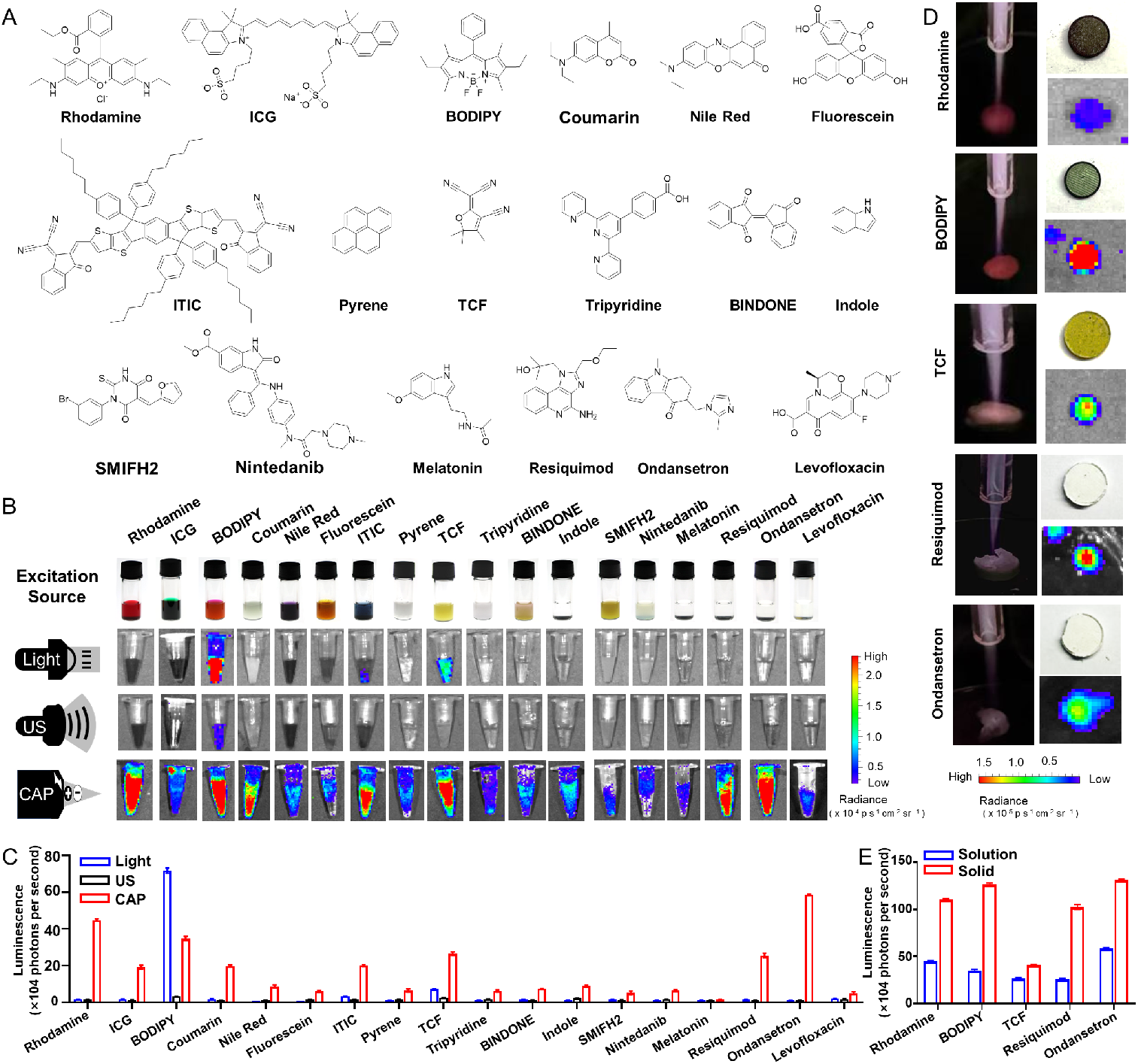
The CAP-induced afterglow luminescence from organic matters. **(A)** The molecular structure of various representative organic molecules. **(B)** The afterglow imaging of the aqueous solution of the organic molecules with different excitation source treatments including white light, ultrasound and CAP. **(C)** The luminescence intensity analysist of the aqueous solution of the organic molecules with different excitation source treatments. **(D)** The afterglow imaging of the solid of the organic molecules with CAP treatment. **(F)** The luminescence intensity analysist of the solid of the organic molecules with different excitation source treatments. All data were obtained from three times of independent experiments and shown as mean ± SD(*p< 0*.*001*).

### Investigation of CAP luminescence imaging

We focused on studying the luminescence performance of rhodamine using CAP induced luminescence imaging mode, which involves imaging after stopping CAP excitation. We excited Rhodamine with CAP at different times (i.e. 30s-5min) and observed similar afterglow spectra. Within the processing time range of 2min-3min, Rhodamine afterglow was most strongly excited (Figure 2A-B). The CAP induced afterglow intensity of Rhodamine shows a positive linear relationship with molecular concentration (Figure 2C-D). After CAP excitation, we observed that the afterglow signal of rhodamine lasted for 2 hours (Figure 2E-F). In addition, we also detected the wavelength range of CAP induced afterglow of Rhodamine, between 600nm-720nm (Figure 2G-H). The fluorescence spectrum peak of Rhodamine was in the range of 600nm-660nm, (Figure 2I-J) which is similar to the afterglow range induced by CAP.

**Figure 2.**
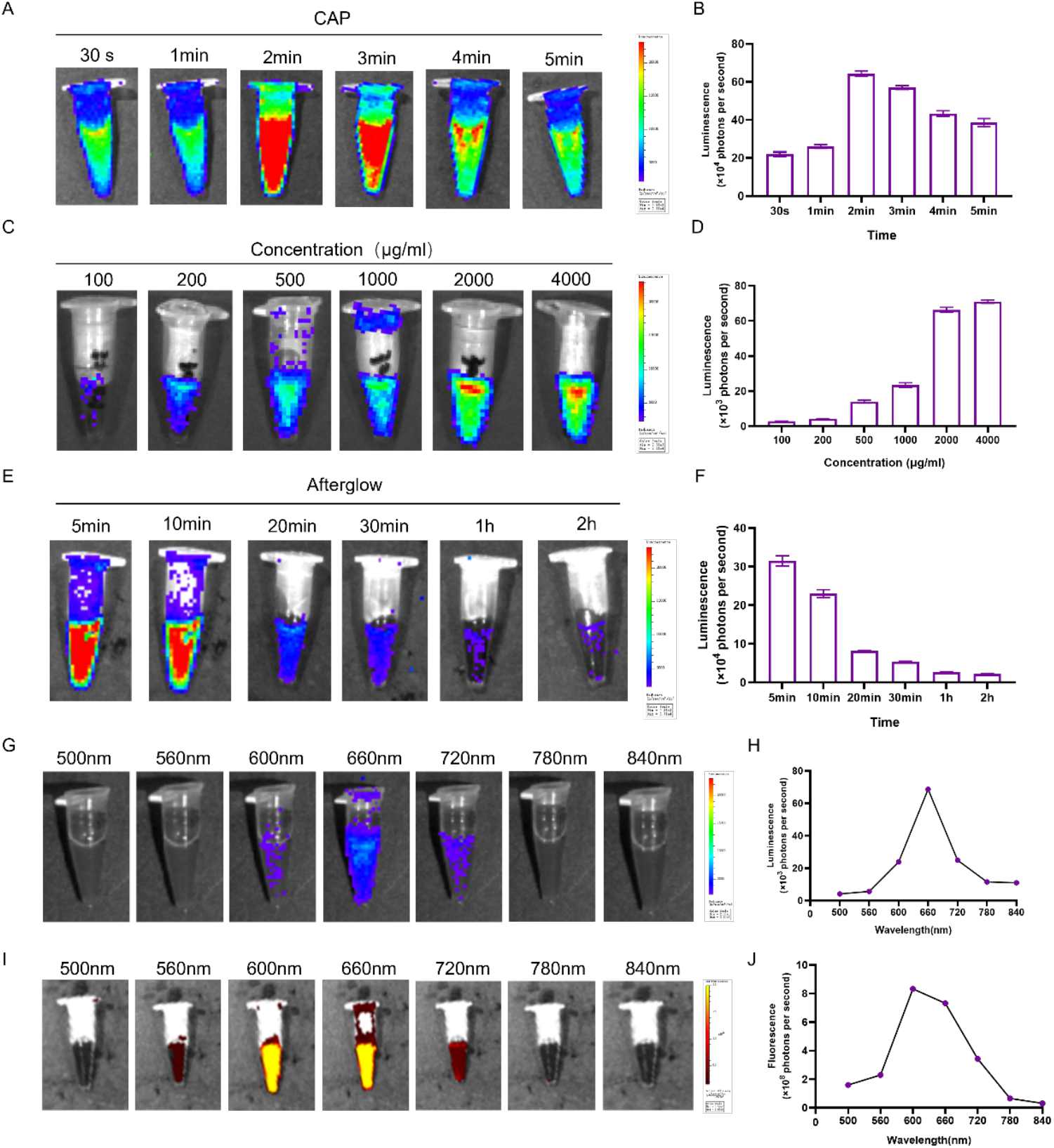
The afterglow luminescence characteristics of Rhodamine material after CAP treatment. **(A-B)** Afterglow luminescence of rhodamine materials treated with cap at different times. **(C-D)** The afterglow luminescence intensity of Rhodamine materials with concentration gradient after CAP treatment. **(E-F)** Continuous afterglow luminescence time of rhodamine material after 3 minutes of cap treatment, **(G-H)** The afterglow luminescence characteristics of Rhodamine materials excited by light with different excitation wavelengths (500-840nm) **(I-J)** Fluorescence luminescence characteristics of Rhodamine materials excited by light with different excitation wavelengths (500-840nm). All data were obtained from three times of independent experiments and shown as mean ± SD(*p<0*.*001*).

### Mechanism of CAP induce afterglow luminescence imaging

Optical emission spectroscopy and plasma diagnostics confirmed that the CAP jet simultaneously delivers energetic electrons, reactive oxygen and nitrogen species, weak UV photons, and a localized electric field (Figure 3A-B). These multi-modal energy inputs likely act synergistically to populate carrier traps and/or generate new surface states that facilitate long-lived emission. Importantly, clinically relevant compounds, including several FDA-approved drugs and investigational molecules, showed robust afterglow, indicating that this approach does not require custom synthesis of specialized luminescent materials and can be directly applied to pharmacologically relevant agents. To ensure that CAP activation does not induce structural modification or degradation of the materials, we performed high-resolution mass spectrometry (HRMS) before and after CAP exposure on 5 representative compounds (Figure 3C, Supplementary Figure 1). The spectra remained identical within experimental error, confirming that CAP charging is a physical process that preserves the chemical integrity of the molecules, an essential consideration for clinical translation and safety. It is worth noting that a few new peaks are observed at specific molecular weight positions following the cleavage of the carbon-carbon double bond. This indicates that a negligible amount of new compounds may be generated after CAP treatment, which contributes to the occurrence of afterglow (Supplementary Figure 2). To further identify the critical active component of CAP responsible for inducing afterglow, we performed exclusion experiments and single-factor treatment assays.

**Figure 3.**
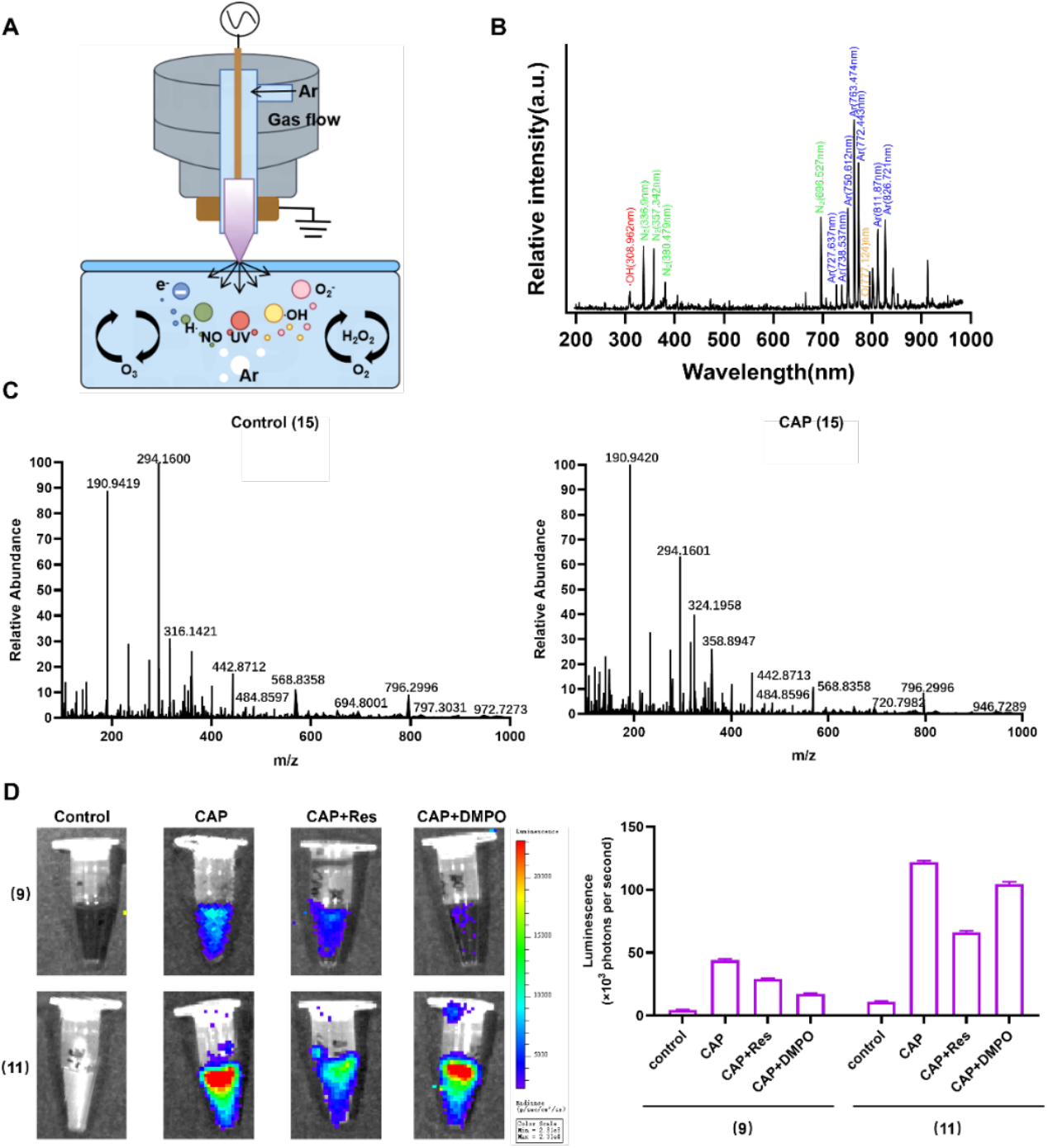
Mechanism of CAP induce afterglow luminescence imaging. (A) Schematic diagram of active substances and UV components contained in CAP. (B) Opticaemission spectrum (OES) of CAP. (C) Mass spectra of material (15) before and after CAP treatment. (D) Afterglow imaging of CAP active substance captured using resorcinol and DMPO. All data were obtained from three times of independent experiments and shown as mean ± SD(*p<0*.*001*.

Previous experimental results have shown that CAP can only be excited with afterglow when it comes into direct contact with the test material in solution or solid. And when CAP is isolated from the test material in the solution by a centrifuge tube, it cannot be excited to emit afterglow (Supplementary Figure 3-4). This indicates that the ultraviolet radiation in CAP are not effective components for exciting afterglow. Due to the presence of reactive species such as ROS and RNS in CAP, we used resorcinol and DMPO to capture reactive oxygen species. A portion of the test materials showed a decrease in afterglow intensity after being captured by resorcinol free radicals, while another portion of the materials showed a decrease in afterglow intensity after being captured by DMPO free radicals (Figure 3D). The reactive oxygen species in CAP may be the key active ingredients that stimulate the afterglow of the test material.

### Application of CAP luminescence imaging in postoperative recurrence model

In order to validate the in vivo afterglow effect of CAP-induced materials, we utilized CAP to stimulate rhodamine for afterglow generation, which served as the photodynamic stimulation source. We employed the photosensitizer 5-aminolevulinic acid (5-ALA) for intraoperative tumor resection guidance, adopted hydrogel F127 to fill the wound with the aforementioned material, and exploited the afterglow to activate 5-ALA for photodynamic effect generation. This approach enables continuous elimination of residual intraoperative unresectable tumors and prevention of postoperative recurrence (Figure 4A). In vivo validation confirmed that CAP-induced rhodamine is capable of emitting prominent afterglow signals at the tumor site, with sustained luminescence lasting for 1 hour (Figure 4B-C). Tumor surgical navigation was achieved via in situ injection of 5-aminolevulinic acid (5-ALA) (Figure 4D). Following tumor resection, CAP-treated rhodamine loaded into hydrogel F127 was used to fill the surgical wound, and the afterglow was exploited to activate the photodynamic effect. This approach significantly inhibited tumor recurrence with no significant impact on animal body weight (Figure 3E-F). These findings collectively demonstrate the feasibility and safety of the CAP-induced afterglow-mediated photodynamic strategy for preventing postoperative tumor recurrence.

**Figure 4.**
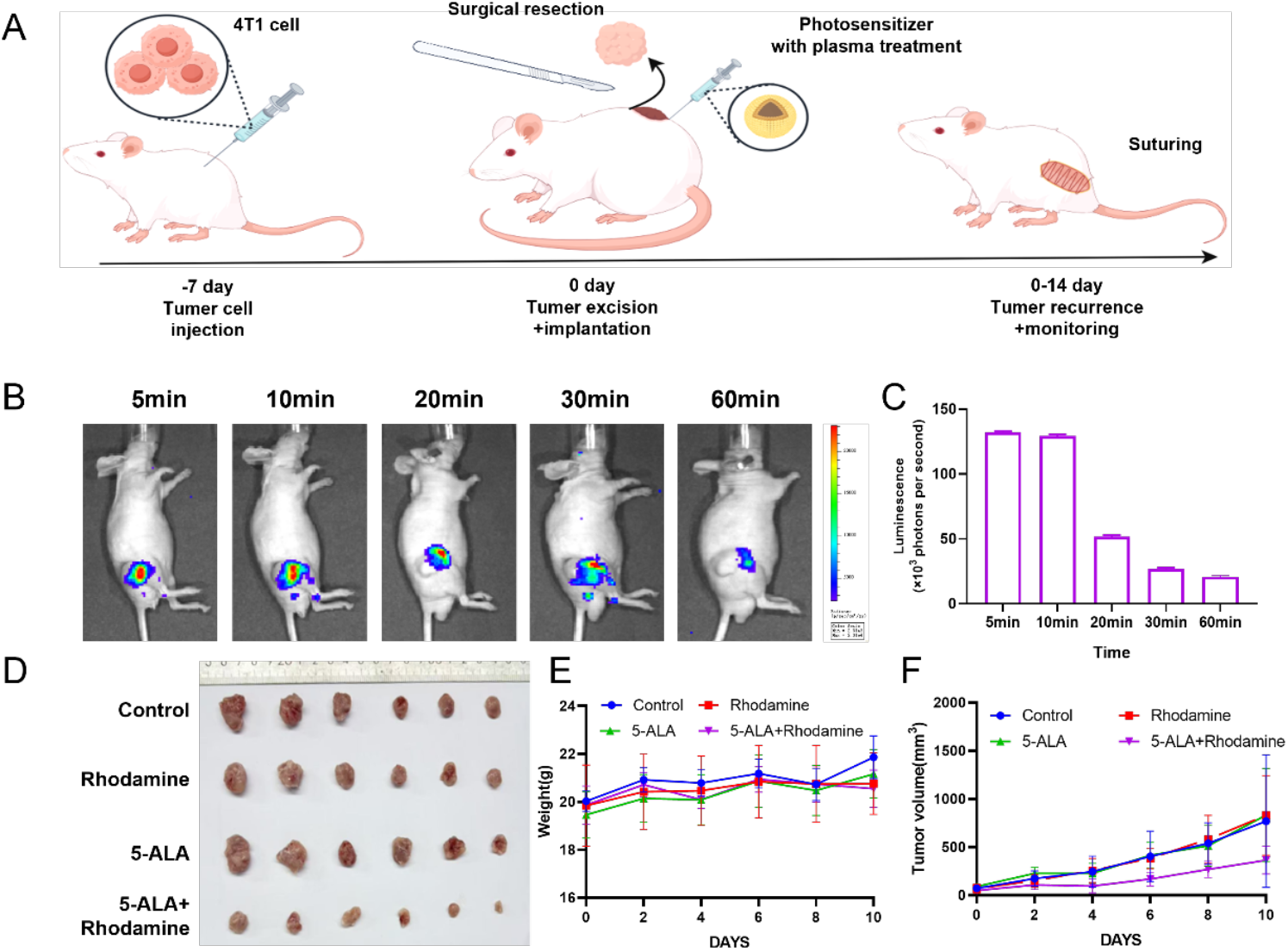
Clinical application of CAP as an excitation source for afterglow materials. (A) Schematic diagram of tumor recurrence model. (B-C) Afterglow imaging of materials treated with subcutaneous injection of CAP in nude mice. (D) Representative macroscopic appearance of tumors after 14 days of subcutaneous injection administration of photosensitizers. (E) The body weight curve in the Oral and Situ model (n= 6). (F) The tumor volume growth curve in the Oral and Situ model (n = 6 *P < 0.05, **P < 0.01).

## Discussion

The discovery that CAP serves as a universal and non-invasive activator of persistent luminescence represents a paradigm shift in the field of stimulus-responsive luminescent materials. This study systematically evaluated 174 compounds and demonstrated that over 32% exhibit detectable afterglow following CAP treatment, with nearly all 20 representative compounds (encompassing clinical drugs, fluorescent dyes, and optoelectronic materials) producing bright, sustained afterglow upon brief CAP exposure (≤3 min). Notably, CAP-induced luminescence intensities were significantly higher than those triggered by conventional light or ultrasound stimulation (p<0.001), overcoming the inherent limitations of traditional excitation modalities that rely on specific photophysical or phononic properties of materials. This universality is particularly striking, as it eliminates the need for custom synthesis of specialized luminescent agents and expands the pool of candidate materials to pharmacologically relevant compounds, including FDA-approved drugs.

The enhanced CAP-induced afterglow in the solid state (3–5 times higher than in solution) provides critical insights into the underlying energy conversion process. The dense and ordered molecular/particle packing in solids minimizes scattering and loss of CAP-excited energy, facilitating efficient conversion of excitation energy into afterglow emission. This observation not only guides the rational design of CAP-responsive luminescent systems but also broadens potential applications, as many practical scenarios (e.g., biomedical devices, sensors) require solid-state materials. For rhodamine, a model compound, the optimal CAP excitation duration (2–3 min), positive linear correlation between afterglow intensity and molecular concentration, 2-hour sustained luminescence, and overlapping afterglow (600–720 nm) with its fluorescence spectrum (600–660 nm) further support the utility of CAP-induced luminescence for quantitative and long-term imaging applications. The consistent spectral profiles across different excitation times also indicate the stability of CAP-induced luminescent processes, a key advantage for reproducible experimental and clinical use.

Mechanistic investigations reveal that CAP’s multi-modal energy inputs (energetic electrons, reactive oxygen and nitrogen species [ROS/RNS], weak UV photons, and localized electric fields) act synergistically to populate carrier traps and/or generate new surface states, driving long-lived emission. Crucially, high-resolution mass spectrometry (HRMS) confirmed that CAP activation preserves the chemical integrity of molecules, with only negligible formation of new compounds via carbon-carbon double bond cleavage—an essential prerequisite for clinical translation and safety. Exclusion experiments and single-factor assays further identified ROS as the critical active component of CAP, as ROS standards (but not RNS standards) induced afterglow in test materials, and ROS scavengers (resorcinol and DMPO) attenuated afterglow intensity. This mechanistic clarity not only advances our understanding of plasma-material interactions but also enables targeted optimization of CAP parameters to enhance luminescence efficiency for specific applications.

The in vivo validation in a postoperative tumor recurrence model highlights the translational potential of CAP-induced afterglow-mediated photodynamic therapy. By combining CAP-stimulated rhodamine (as a sustained afterglow source), 5-aminolevulinic acid (5-ALA, a clinical photosensitizer for intraoperative navigation), and F127 hydrogel (for localized delivery), we achieved continuous elimination of residual unresectable tumors and significant inhibition of recurrence without compromising animal body weight. This strategy addresses a major clinical unmet need: the inability of conventional intraoperative imaging and adjuvant therapies to fully eradicate microscopic residual tumor cells, which are the primary drivers of postoperative recurrence. The 1-hour sustained in vivo afterglow at the tumor site ensures prolonged activation of photodynamic therapy, while the biocompatibility of CAP and the use of clinically approved components (5-ALA, F127) minimize safety concerns and accelerate translational progress.

Despite these promising findings, several limitations warrant further exploration. First, while 32% of tested compounds exhibited CAP-induced afterglow, the structural and chemical features that determine CAP responsiveness remain to be fully elucidated—future studies should focus on structure-activity relationship analyses to predict and optimize candidate materials. Second, the specific ROS species (e.g., singlet oxygen, hydroxyl radicals) responsible for afterglow induction require precise identification to refine activation strategies. Third, the in vivo efficacy and safety need to be validated in larger animal models and different tumor types to confirm clinical applicability. Additionally, optimizing CAP parameters (e.g., treatment duration, intensity) and delivery systems (e.g., minimally invasive CAP probes) could further enhance luminescence intensity, duration, and targeting accuracy.

## Conclusion

This study establishes CAP as a universal, non-invasive, and safe activator of persistent luminescence, with unique advantages over conventional stimuli. The mechanistic insights into ROS-mediated luminescence induction, combined with the successful in vivo application for preventing postoperative tumor recurrence, open new avenues for the development of innovative imaging and therapeutic strategies. Beyond oncology, CAP-induced luminescence could find broad utility in areas such as biosensing, drug delivery, and optoelectronics, where universal and non-destructive excitation of luminescence is desirable. This work not only advances fundamental knowledge of plasma-material interactions but also lays the foundation for translational research that bridges plasma physics, materials science, and biomedicine.

